# Window to the Fluorescence: The *White* Eye-Color Gene of Western Corn Rootworm, *Diabrotica virgifera virgifera*

**DOI:** 10.1101/552935

**Authors:** Nathaniel Grubbs, Fu-Chyun Chu, Marcé D. Lorenzen

## Abstract

Eye-color mutations have proven useful in multiple insect species to help facilitate the development and use of transgenic tools for functional genomics. While there is species-specific variation in the pigments used to color insect eyes, every species studied thus far requires an ortholog of the ABC transporter gene *white* for proper pigmentation of the eyes. Previously, we generated transgenic western corn rootworm, *Diabrotica virgifera virgifera*, and found that their wild-type eye color obscured our ability to visualize a fluorescent marker driven by the widely used 3xP3 eye-specific promoter. Therefore, we sought to identify the *D. v. virgifera* ortholog of *white* (*Dvvw*). Here we report the discovery, cloning, and analysis of *Dvvw* cDNA and promoter. We also utilize RNA interference to knock down *Dvvw* mRNA in a transgenic strain, thereby demonstrating the utility of eye-color mutations when developing transgenic technologies.

## INTRODUCTION

The western corn rootworm, *Diabrotica virgifera virgifera*, is a major pest of corn crops in the US and Europe. Moreover, this beetle is highly adaptable, rapidly developing resistance to a number of control methods, including chemical controls and crop rotations (Reviewed in (Miller *et al*. 2009)). New innovations in pest control, such as transgenic corn expressing beetle-specific toxins, have helped farmers maintain productivity (Reviewed in (Gray *et al*. 2009)), but understanding of the genetic sources of adaptability lag far behind *D. v. virgifera*’s ability to overcome new controls. One reason for this may be its large genome, ~2.58 Gbp which is more similar in size to that of humans and over 10x the size of the genomes of insect model organisms, such as the fruit fly, *Drosophila melanogaster*, and the red flour beetle, *Tribolium castaneum* (Adams *et al*. 2000; Sappington *et al*. 2006; Richards *et al*. 2008; Coates *et al*. 2012). Interestingly, this genome appears to have been enlarged, at least in part, by the expansion of repetitive sequences, thus increasing the difficulty of genome assembly (Coates *et al*. 2012).

Without a fully sequenced and assembled genome, reverse genetic studies are more difficult to carry out. For example, RNA interference (RNAi) is highly effective in *D. v. virgifera*, and acts systemically to knock down target gene expression (Baum *et al*. 2007; Alves *et al*. 2010), allowing its use for functional studies in this species. However, using RNAi to study gene function requires knowing the sequence of the target. While transcriptomic data can fill this gap, RNAi is still limited to predictable targets, biasing research away from novel or poorly studied genes, which would be the most likely types of genes to contribute to *D. v. virgifera*’s adaptability. Transgenic technologies could be used to overcome these limitations with methods like the Jumpstarter mutagenesis system (Horn *et al*. 2003; Lorenzen *et al*. 2007), which disrupts gene function through random hopping of trackable transgenes.

Like RNAi, genome editing using RNA-guided nucleases, such as Cas9, have proven to be easily deployed in a number of species. However, the cutting efficiency of Cas9 is dependent on a number of variables, resulting in the need for some trial and error for each genomic locus researchers wish to target. Efficiency of genome editing can be improved with cellular expression of Cas9 (Ren *et al*. 2013), which is possible with traditional transgenic tools. However, these and other transgenic functional genomics techniques depend on reliable transformation markers.

Since the early days of transposon-mediated germline transformation in *D. melanogaster*, eye-color genes have been the gold standard for transformation markers (Rubin and Spradling 1982; White *et al*. 1996; Besansky *et al*. 1997; Sumitani *et al*. 2005). Indeed, even with the advent of fluorescent markers, eye-specific expression has been favored over other promoters, via the development of the 3xP3 artificial eye-specific promoter, since it is typically easy to identify (Berghammer *et al*. 1999; Horn *et al*. 2000). The limited scope of expression also reduces concerns of toxicity, while enabling researchers to more readily distinguish transformed individuals from general autofluorescence (Berghammer *et al*. 1999; Horn *et al*. 2002).

More importantly, the 3xP3 promoter appears to function almost universally, producing eye expression in many of the species tested (Berghammer *et al*. 1999; Kokoza *et al*. 2001; Gonzalez-Estevez *et al*. 2003). In addition, it does not seem to interfere with enhancer traps or other site-specific enhancements of marker gene expression, allowing marked constructs to be used for tagging new genes and enhancer regions (Horn *et al*. 2002). These advantages were exploited when a Jumpstarter mutagenesis system was being developed in *T. castaneum* (Lorenzen *et al*. 2007); a line carrying a muscle-specific enhancer trap was used as a donor, and successful jumps were confirmed by the loss of muscle, but not eye-specific, EGFP expression.

In spite of the many advantages of eye-specific markers, the use of eye-expression markers is limited by the availability of eye-color mutants. Even when using 3xP3-driven fluorescent markers, integrations into expression-quenching genomic locations can be further masked in individuals with fully-pigmented eyes (Horn *et al*. 2000; Lorenzen *et al*. 2002), making eye-color mutants a useful lab tool. For this reason, we have sought to knockout eye-color production in *D. v. virgifera*.

Although many of the insect species that have been studied produce only a single type of eye pigment, ommochromes (Dustmann 1968; Beard *et al*. 1995; Cornel *et al*. 1997; Quan *et al*. 2002; Moraes *et al*. 2005), some species, including *D. melanogaster*, employ an additional pigment type (Summers *et al*. 1982). In either case, total loss of function in the orthologs of the *D. melanogaster* ABC transporter, White, result in total loss of visible eye pigments (Sullivan and Sullivan 1975; Sullivan *et al*. 1979; Besansky *et al*. 1995; Abraham *et al*. 2000; Sumitani *et al*. 2005; Grubbs *et al*. 2015). Since we do not know the nature of western corn rootworm eye pigments, we decided to seek out and target the *D. v. virgifera white* (*Dvvw*) ortholog to ensure we would significantly impact eye-color development. Here, we report the identification of *Dvvw*, and demonstrate its function by RNAi-mediated gene silencing.

## MATERIALS AND METHODS

### Insects and egg collection

The non-diapausing strain of *D. v. virgifera* (Kim et al., 2007) used in this experiment is a mixed colony composed of otherwise wild-type individuals received from Dr. Wade French (USDA-ARS-NGIRL, Brookings, SD), and Crop Characteristics, Inc. (Farmington, MN, USA). The strain expressing red fluorescence (DsRed) in their eyes are the Min-2 strain developed by Chu *et al*. (2017). Colony size was maintained at >500 individuals per 30 cm^3^ cage (BugDorm, MegaView Science, Taiwan) to minimize the effects of inbreeding. Adults were reared on artificial diet (Product# F9766B; Frontier Agricultural Services, Newark, Delaware, USA) and held at 26°C, 60% humidity with a 14:10 h (light:dark) photoperiod. Beetles were reared essentially as in (Georce and Ortman 1965; Branson *et al*. 1988). Egg collections were carried out essentially as in (Chu *et al*. 2018).

### Cloning *Dvvw* coding sequence and promoter

The sequence of the *T. castaneum* White protein was used as query against an in-house database consisting of transcriptomic data from mixed-stage *D. v. virgifera* embryos (~500 eggs from an overnight egg lay aged up to 14 days), mixed-stage larvae (20 first-instar larvae; 10 second-instar larvae; and 2 third-instar larvae), as well as an adult male, and an adult female (MiSeq, Illumina, San Diego, CA, USA), resulting in the identification of an EST contig with high identity to the query. This data was used to design the primers, Dvv-w FL F2 and Dvv-w FL R1 (all primer sequences can be found in Supplemental Table 1), for amplifying the full-length coding sequence. The template for cloning was derived from total RNA isolated from mixed-stage *D. v. virgifera* embryos (~500 eggs from an overnight egg lay aged up to 14 days) using an RNeasy^®^ Mini Kit (QIAGEN, Valencia, CA, USA) following the manufacturer’s instructions. This RNA was converted to cDNA using an oligo dT primer (RT-Uni) and Superscript III (ThermoFisher Scientific, Waltham, MA, USA) for use as a PCR template. The resulting amplicon (1.8 kb) was ligated into TOPO™ TA PCR4 vector (ThermoFisher), and the resulting clone sequenced (GENBANK accession no. XXXXXX). The *Dvvw* gene structure was then determined by querying (BlastN) the *D. v. virgifera* whole genome sequence contigs database (WGS project ID PXJM) at NCBI. This yielded two scaffolds which were downloaded into VectorNTI (Invitrogen, Carlsbad, CA, USA) so we could determine exon and intron lengths.

Genomic DNA (gDNA) was isolated from four adults (~25 mg of tissue) using a DNeasy^®^ Blood & Tissue kit (QIAGEN), following the manufacturer’s Animal Tissues protocol. For the *Dvvw* promoter, approximately 25 ng of gDNA and the Dvv-w 5prime F1 and Dvv-w 5prime R primers were used to amplify a 1069-bp portion of the genome (including the first exon and 927 bases of upstream sequence), which was cloned using the pGEM^®^-T Easy Vector System (Promega, Madison, WI, USA), and the resulting clone sequenced (GENBANK accession no. XXXXXX). The *D. v. virgifera aspartate 1-decarboxylase* (*ADC*) gene was used as a control. For *DvvADC* dsRNA, a 621-bp fragment was amplified from 25 ng of gDNA using first-round primers. Second-round primers, bearing a T7 RNA polymerase promoter sequence, were then used to amplify a 387-bp fragment which was then used to generate double-stranded RNA as described below.

### RNAi and microinjection

DNA templates were amplified using iProof™ proof reader polymerase (BioRad, Hercules, CA, USA) (T7 and M13R-T7 primers were used for *Dvvw*). These PCR products were purified using a QIAquick PCR Purification Kit (QIAGEN, Hilden, Germany), then 1 μg of each was used as template for synthesis of dsRNA using a MEGAscript™ T7 Transcription Kit (ThermoFisher). The MEGAclear™ Transcription Clean-up Kit (ThermoFisher) was used to purify the dsRNA, which was eluted into nuclease-free water.

For microinjection, the final concentration of *Dvvw* dsRNA was brought to 1000 ng/μl and 20% Phenol Red (Cat #143-74-8; Sigma-Aldrich, St. Louis, MO, USA) added for coloration. We injected late stage *D. v. virgifera* third-instar larvae as summarized in Table 1. All injectees were reared to adulthood in a container of soil with sprouted corn, then screened for eye-color phenocopy.

**Table 1:** *Dvvw* RNAi results

| Solution | Strain | # Inj Larvae | # Adults | # w/ Eye Phenotype |
| --- | --- | --- | --- | --- |
| Buffer | WT | 10 | 4 | 0 |
| *Dvww* dsRNA | WT | 22 | 6* | 4 |
| *Dvww* dsRNA | Min-2 | 20 | 9 | 4 |
| *DvvADC* dsRNA | WT | 12 | 5 | 0 |
| *DvvADC* dsRNA | WT | 21 | 15 | 0 |

\*11 originally survived, but 5 died before they could be screened.

### White ortholog identification, protein alignments and phylogeny

White orthologs and the two Scarlet sequences from other insect species had previously been identified for (Grubbs *et al*. 2015). The Maximum Likelihood phylogenetic tree was constructed in the MEGA X 64-bit program, version 10.0.5 (Kumar *et al*. 2018), using default parameters in all categories except: bootstrapping with 500 replicates, LG model of amino-acid substitution with Gamma distributed substitution rates with Invariant sites (based on Best Model determination within the MEGA program), and Partial Deletion treatment of gaps/missing data (Hall 2013). Figure S1 was generated using an alignment constructed by the PSI-Coffee tool at tcoffee.crg.cat (Notredame *et al*. 2000; Tommaso *et al*. 2011).

## RESULTS

ABC transporters responsible for importing eye pigments in insects typically possess a very specific motif (CDEPT) within their Walker B ATP-binding domain (Grubbs *et al*. 2015). To identify *Dvvw*, we queried (TBlastN) a *D. v. virgifera* RNA-Seq database with the deduced amino-acid sequence of the *T. castaneum white* (*Tcw*) gene, and were able to identify a contig encoding the expected CDEPT motif. The predicted translation product of this contig was otherwise similar to TcW, so we confirmed its expression in *D. v. virgifera* embryos by amplifying a nearly full-length transcript from embryonic cDNA via PCR. The longest 5’ UTR is 115 bases in length, the CDS contains 1974 bases encoding a protein 657 amino acids long, and the 3’ UTR is 154 bases. The *Dvvw* 3’ UTR is much longer than its *T. castaneum* ortholog; otherwise, their features are comparable in length (Grubbs *et al*. 2015).

The deduced amino-acid sequence of *Dvvw* shares 59% identity with its *T. castaneum* counterpart, and 52% identity with *D. melanogaster* W (DmW), which is comparable to the 53% identity TcW shares with DmW. Phylogenetic analysis shows that DvvW is distinct from the Scarlet (St) lineage (Figure 2), reinforcing our conclusion that this is the W ortholog. As expected, the two beetle W orthologs, DvvW and TcW, form a distinct group, as do the Dipteran orthologs, separating from the remaining species included in our analysis (*Bombyx mori* and *Apis mellifera*). Both beetle proteins are notably shorter than those of the other species analyzed, with most of the difference in length found in the highly-variable N-terminus (Figure S1). Otherwise, the proteins are very consistent, and almost invariable in critical domains.

**Figure 1:**
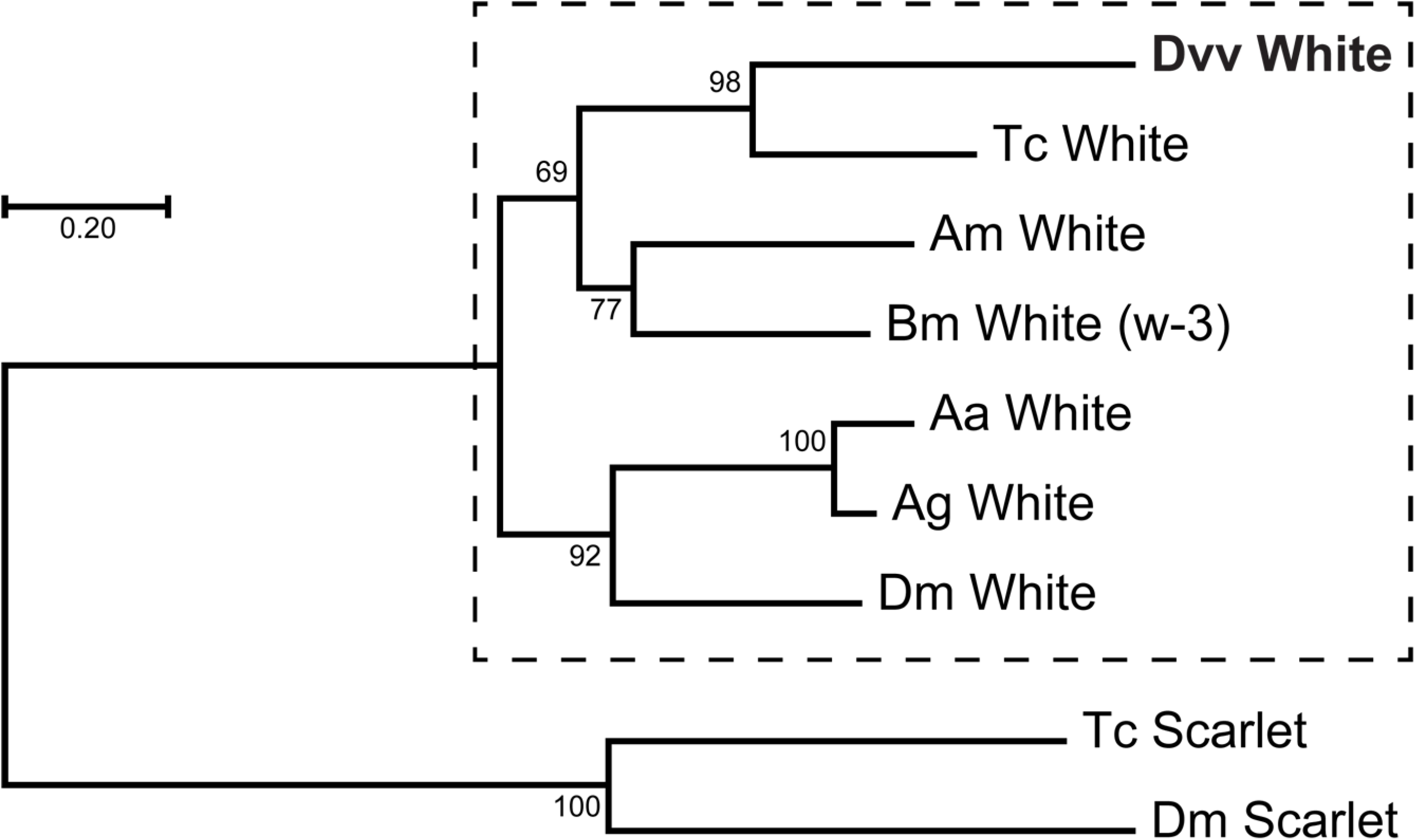
Phylogeny of insect White orthologs. *D. v. virgifera* White (bold) groups with other White orthologs (dashed box), not with Scarlet orthologs, which were used as an outgroup to root the tree. Aa: *Aedes aegypti*, White AAEL016999-PA; Ag: *Anopheles gambia*, White AGAP000553-PA; Am: *Apis melifera*, White modified from XP_026299500.1; Bm: *Bombyx mori*, White NP_001037034.1; Dm: *Drosophila melanogaster*, White NP_476787.1, Scarlet NP_524108.1; Tc: *Tribolium castaneum*, White NP_001034521.1, Scarlet NP_001306193.1.

**Figure 2:**
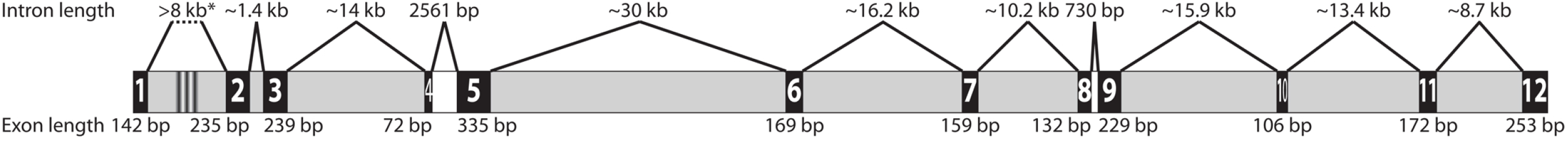
*Dvvw* gene structure. Exons are in black, numbered, and to scale with length listed below. Grey spaces indicate introns that span one or more gaps between scaffold 02243 contigs; white introns lie between exons on the same contig. The black stripes in the first intron indicate the transition between scaffolds. Introns are 1:10 scale relative to exons, with lengths listed above.

Using the complete *Dvvw* cDNA sequence, we were able to query the *D. v. virgifera* genome available on NCBI (GENBANK accession no. GCA_003013835.2) with BlastN to identify the exon boundaries and arrangements (Figure 2). This revealed twelve exons across ten separate contigs on two scaffolds. Exon 1, which contains the 5’ UTR and the first nine codons of the CDS, is the only exon on Dvir_v2_scaffold_02154 (GENBANK accession no. ML017136.1), while the remaining exons are found on Dvir_v2_scaffold_02243 (GENBANK accession no. ML017225.1). *Dvvw* spans more than 123 kb, from the end of scaffold 02243, 5’ of exon 2, to the 3’ end of exon 12. The length and number of the exons is not unusual for an eye color ABC gene, but the length of the introns is significantly increased relative to organisms like *T. castaneum*. The longest intron appears to be between exons 5 and 6, which is at least 30 kb, but may be longer, depending on the actual sizes of the gaps between intervening contigs. The distance between exons 1 and 2 may be greater still, but cannot be reliably deduced, since they are on separate scaffolds. However, the distance between the 3’ end of exon 1 and the end of scaffold 02154 is more than 45 kb, so if there is no overlap between these two scaffolds, exons 1 and 2 could be separated by more than 53 kb. Only two pairs of exons share a contig: exons 8 and 9 on contig 42, which flank the shortest intron of 730 bp, and exons 4 and 5 on contig 48, which flank the third shortest intron of 2561 bp. Unlike the introns, the exons of *Dvvw* are all of similar length. Exons 4 and 5 happen to be the shortest (72 bp) and longest (335 bp) exons, respectively.

Since *Dmw* and *Tcw* are each expressed from TATA-less promoters (Grubbs *et al*. 2015), we sought to identify promoter sequences that may drive expression of *Dvvw* for comparison. To accomplish this, we cloned ~1-kb upstream of the *Dvvw* translation start site for sequencing (−927 to +142, where +1 corresponds to the 5’-most nucleotide of the longest *Tcw* cDNA, Figure 3). Analysis of the sequencing results revealed two putative promoters (score = 0.98 and 0.83, 5’-3’, Neural Network Promoter Prediction). Surprisingly, both of these promoters appear to contain canonical TATA boxes (Ohler *et al*. 2002). In addition, two extended TATA motifs (FitzGerald *et al*. 2006) were found elsewhere in this region, one upstream and one downstream of the putative promoters. Two potential initiators (INR), similar to the extended motifs defined by Ohler et al. (2002), were identified; one sits at the end of the first putative promoter at a canonical distance from the TATA box (Juven-Gershon *et al*. 2008), while the other lies between positions +7 and +20, near the start of transcription, as would be expected. Two canonical downstream promoter elements (DPE) are also present (Ohler *et al*. 2002), but the first lies at the start of the second putative promoter, while the other is upstream of the start of transcription (−73 to −67). Several other downstream elements (Arkhipova 1995) can be found throughout the region (Figure 3), with a particular enrichment of downstream triplets and tetramers within the 5’UTR. No other promoter motifs (Ohler *et al*. 2002; FitzGerald *et al*. 2006) were found in the cloned sequence, although the 5’ UTR possesses false start codons, as was seen in *Tcw* (Grubbs *et al*. 2015); translation from these sites would terminate quickly, making at most a protein of 43 amino acids in length.

**Figure 3:**
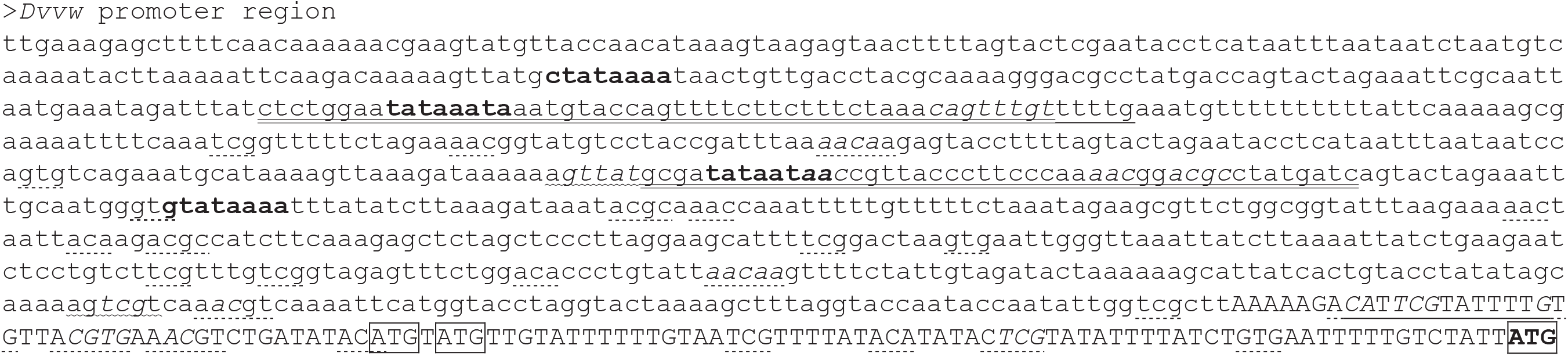
Nucleotide sequence of the putative *Dvvw* promoter region. Capital letters indicate the longest transcript detected (first capital base = +1), with the premature ATGs in boxes, while the boxed and boldfaced ATG indicates the start of the proper coding sequence. Proposed initiator sequences are underlined, clusters of downstream elements are denoted by dashed underlines, potential DPE motifs are marked with squiggly underlines, predicted promoter sequences are double-underlined, and overlapping features are italicized. Boldface type denotes consensus TATA boxes.

To confirm the role of *Dvvw* in eye pigmentation, we performed RNAi by injecting late-stage larvae with dsRNA derived from the cloned *Dvvw* cDNA. Results are summarized in Table 1. Survival rates are low for all injection rounds, including buffer-only controls, ranging from 71% to 40%. However, a reduction of eye pigmentation was only observed in individuals injected with *Dvvw* dsRNA. Pigment levels did not change through the course of observations, which were carried out over one week post-eclosion. Just as in *T. castaneum*, reduced *w* function in *D. v. virgifera* had no impact on the ocular diaphragm, leaving a ring of dark pigment around the edge of the eyes (Figure 4).

**Figure 4:**
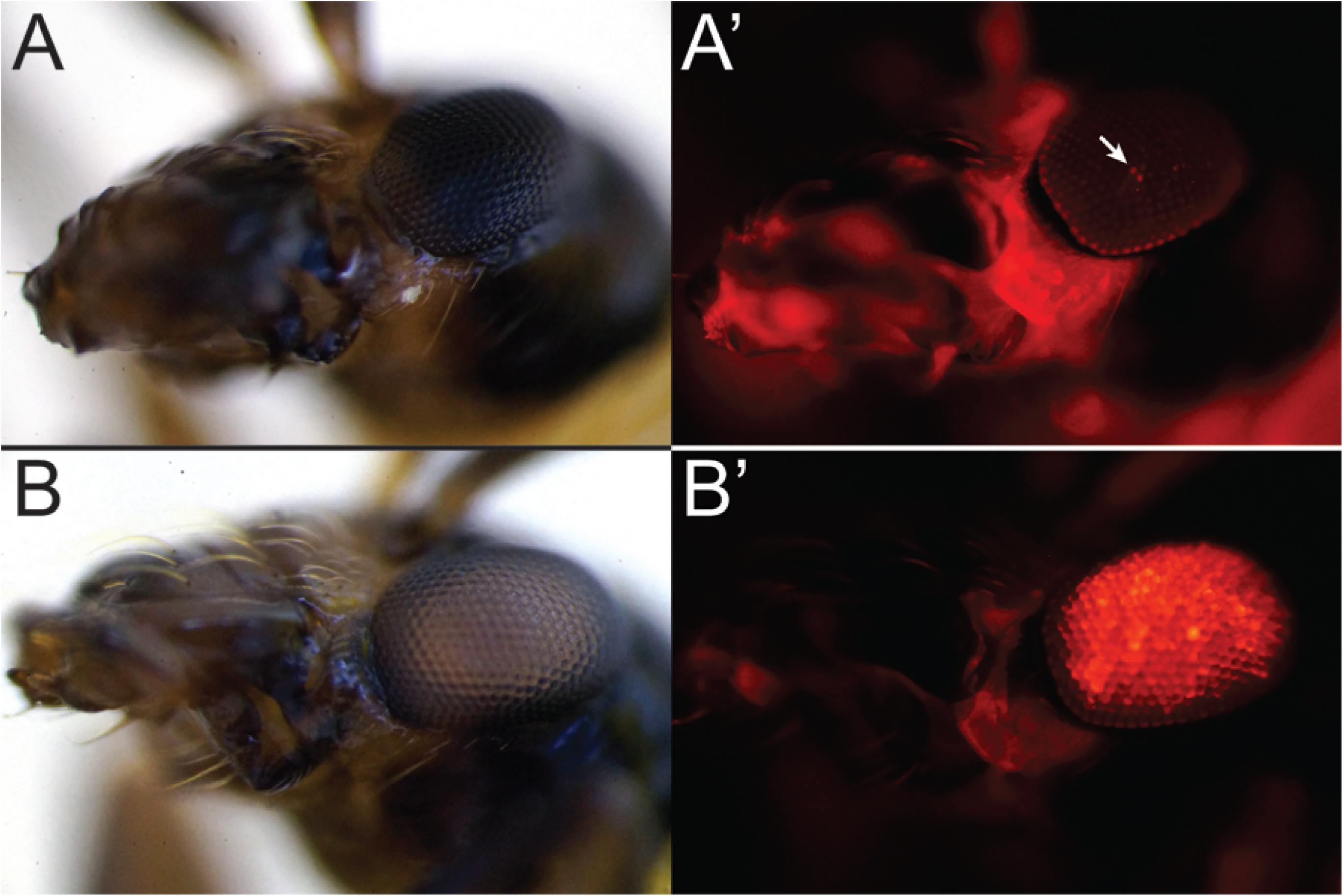
Effect of *Dvvw* RNAi on eye color. Buffer (A) and *Dvvw*-specific dsRNA (B) eye color of Min-2 strain under white light (left) and red fluorescent filter (right). Note that, with wild-type eye color, DsRed fluorescence is only visible in a few ommatidia (arrow, A’), while DsRed fluorescence is clearly visible throughout the eye when pigmentation is reduced (B’).

Since our hypothesis is that reduction of eye pigmentation will improve our ability to visualize eye-specific fluorescent markers, we tested the impact of *Dvvw* RNAi in a transgenic strain. The Min-2 strain developed previously by our lab (Chu *et al*. 2017) carries a transposon marked with 3xP3 driving DsRed. Survival and knockdown rates of *Dvvw* RNAi in Min-2 (Table 1), as well as the phenotype were comparable to those observed in the WT strain. More importantly, while the wild-type eye pigment of *D. v. virgifera* interferes with our ability to visualize the fluorescent red protein (Figure 4A’), RNAi knockdown of *Dvvw* allows the fluorescence to be seen clearly throughout the eye (Figure 4B’).

## DISCUSSION

In this study, we have identified the *D. v. virgifera* ortholog of the eye-pigment transporter, *white*, by RNA-Seq, followed by cloning of the full-length transcript. Using this data, we have been able to identify a potential promoter region for this gene, and to place this gene in a larger evolutionary context. Finally, we have reduced the function of *Dvvw* using RNAi, and have shown that it does, indeed, play a role in proper pigmentation of the adult eye.

Given the findings in other species, it was not surprising to see near-total loss of pigmentation in *D. v. virgifera* eyes as a result of RNAi targeting *Dvvw*. More importantly, this reduction in eye pigmentation is, indeed, helpful in visualizing eye-specific fluorescent markers. Such markers would otherwise be difficult to observe through the wild-type pigmentation, making detection of transgenic lines more difficult, as more time and effort would then need to be spent screening. So, it is clear that targeting the *Dvvw* gene for mutation will accomplish our primary goal – creating a white-eyed background in which we can easily view fluorochromes expressed in the eye, or to serve as a background for eye-color rescue.

Transport of ommochrome pigments into insect eyes requires two ABC transporters, W and its paralog, Scarlet (St) (Tearle *et al*. 1989). In species that only employ ommochromes to color eyes, total loss of St function can resemble loss of W (Tatematsu *et al*. 2011; Grubbs *et al*. 2015). Thus, it was imperative for us to clearly determine the identity of the newly-discovered gene. Our phylogenetic analysis clearly shows that DmSt and TcSt group together, separated from the W orthologs, including DvvW, affirming the identity of our newly-discovered *D. v. virgifera* gene. While identification of *Dvvst* would certainly help solidify this conclusion, we were unable to find sequences matching this gene in our RNA-Seq data. Similarly, *Tcst* does not appear to be expressed in *T. castaneum* embryos, even though *Tcw* is. If these results are consistent, then we can conclude that this particular expression profile is evolutionarily conserved, but more RNA-Seq data is necessary, particularly from different life stages and more species, to determine what shapes these expression patterns, and what their effects on development might be.

Our analysis of the genomic sequence upstream of *Dvvw* revealed some interesting insights into sequence motifs that may drive expression. Surprisingly, unlike the orthologs from *D. melanogaster* and *T. castaneum* (Grubbs *et al*. 2015) the *Dvvw* promoter region appears to contain multiple TATA boxes. More interesting is the prevalence of common motifs. INR, TATA, and DPE were all found in the region at least twice, in addition to a number of other downstream trimer and tetramer motifs. Studies of *D. melanogaster* promoters suggest that while INR motifs often pair with either TATA or DPE, TATA and DPE are not commonly found together (FitzGerald *et al*. 2006). One possibility raised by the numbers and positions of these common promoter elements is that *Dvvw* has multiple promoters, either as false starts, or for temporal and positional control. Indeed, the presence of multiple potential translation start sites in the 5’ UTR of this gene may lend credence to the use of these elements in fine-tuning expression. It is important to note that our RNA-Seq data did not include pupae, a critical stage for the development of adult eye pigmentation. Future work would benefit from a more detailed examination of expression from all life stages, as well as from specific tissues, in order to clarify this question. Unfortunately, what we know of insect promoter function is derived almost exclusively from *D. melanogaster*, so it is also possible that these motifs are used differently in *D. v. virgifera*. More research needs to be done on arthropod promoter regions to determine what features are shared across the group, and what may be species specific. Adding more data from the *D. v. virgifera* transcriptome and genome will help us begin to answer these questions.

*D. v. virgifera* remains an important agricultural pest, and only now are the tools necessary to study it genetically becoming a reality for researchers. But more is still needed. Here, we have taken the first steps, though small, towards building a transposon based mutagenesis system. By identifying *Dvvw*, we are now able to create a mutant line that will serve as a reliable genetic background for screening eye-specific marker genes. With these tools in hand, we will be able to unlock the mysteries of this perplexing pest.

## Acknowledgements

This work was supported by a grant from the Monsanto Corn Rootworm Knowledge Research Program, grant number AG/1005 (to MDL), and start-up funds to MDL from NC State University. FC was supported by grants from Monsanto’s Corn Rootworm Knowledge Research Program (AG/1005) and the National Science Foundation, grant number MCB-1244772 (to MDL). NG was supported in full by a grant from the National Science Foundation (MCB-1244772). The authors declare no competing interests. NG, FC and MDL conceived and designed the experiments; NG and FC performed the experiments and analyzed the results; and NG, FC, and MDL wrote the manuscript. We thank Pei-Shan Wu, Sofia Pinzi, Teresa O’Leary, Lauren Slayton, Wanose Getachew, Shana Mangaldas, Neelan Patel, Sarah Evans, Kathryn Chalmers, and Ashley Liao for their expert assistance in rearing *D. v. virgifera*.

---

**Figure S1:** Full alignment of White orthologs compared in this work. Amino acids in black highlight are seen in that position in more than half of aligned sequences; grey highlight marks similar amino acids. The consensus line shows the most likely amino acid for a given position, with invariable positions marked with a capital letter.

**Table S1:**
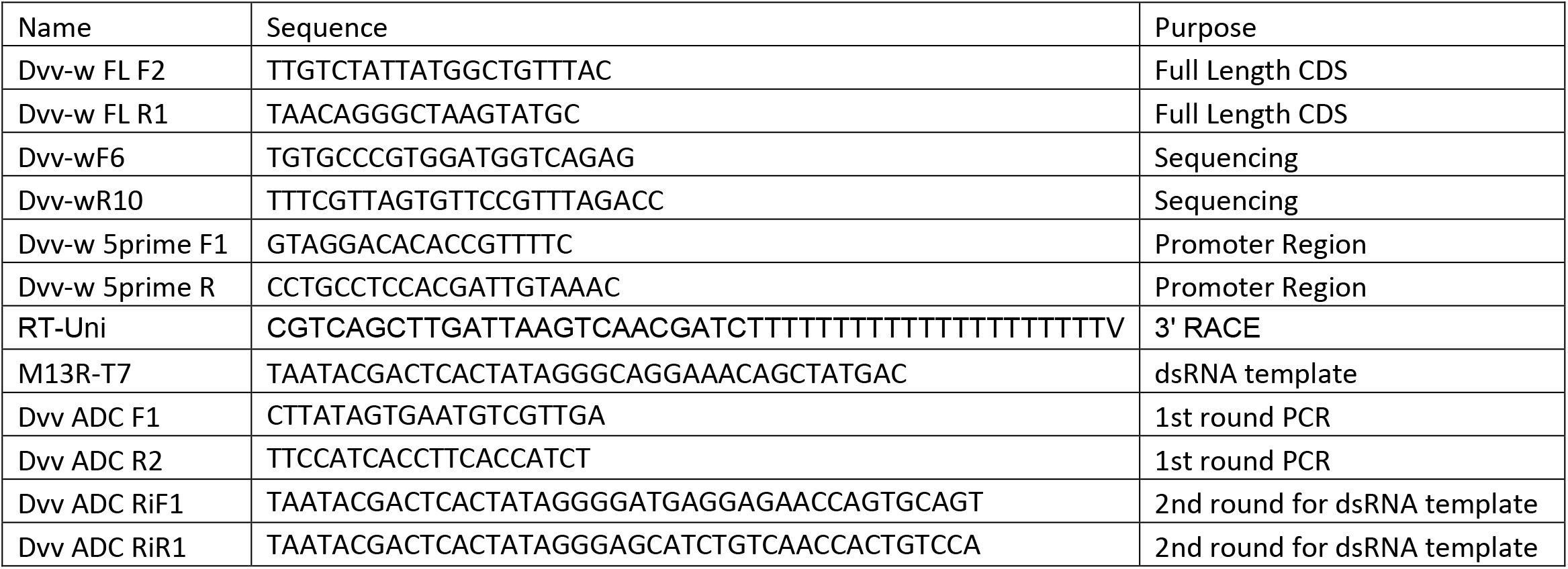
Primers

